# D-amphetamine maintenance therapy reduces cocaine use in female rats

**DOI:** 10.1101/2021.12.08.471850

**Authors:** Ndeye Aissatou Ndiaye, Florence Allain, Anne-Noël Samaha

## Abstract

Currently, there are no approved medications to treat cocaine addiction. In this context, d-amphetamine maintenance therapy is a promising pharmacological strategy to reduce cocaine use. In both male rats and human cocaine users, d-amphetamine treatment reduces cocaine taking and seeking. However, this has not been examined systematically in female animals, even though cocaine addiction afflicts both women and men, and the sexes can differ in their response to cocaine. Here, we determined how d-amphetamine maintenance therapy during cocaine self-administration influences cocaine use in female rats. In experiment 1, two groups of female rats received 14 intermittent access (IntA) cocaine self-administration sessions. One group received concomitant d-amphetamine maintenance treatment (COC + A rats; 5 mg/kg/day, via minipump), the other group did not (COC rats). After discontinuing d-amphetamine treatment, we measured responding for cocaine under a progressive ratio schedule, responding under extinction and cocaine-primed reinstatement of drug seeking. In experiment 2, we assessed the effects of d-amphetamine maintenance on these measures in already cocaine-experienced rats. To this end, rats first received 14 IntA cocaine self-administration sessions without d-amphetamine. They then received 14 more sessions now either with (COC/COC + A rats) or without (COC/COC rats) concomitant d-amphetamine treatment. In both experiments, d-amphetamine treatment did not influence cocaine-primed reinstatement of cocaine seeking. However, after d-amphetamine treatment, rats responded less for cocaine both under progressive ratio and extinction conditions. Thus, d-amphetamine treatment can both prevent and reverse increases in the motivation to take and seek cocaine in female animals.

## 1. Introduction

Across the globe, many people addicted to cocaine seek treatment, but the prevalence of cocaine addiction has not decreased overtime (CTADS, 2017; Administration & Services, 2020). A promising treatment approach involves giving a dopamine agonist with similar pharmacological properties as cocaine, but in a continuous, slow-acting formulation to reduce abuse potential (Kampman, 2019; Negus & Henningfield, 2015; Tardelli et al., 2020). D-amphetamine, a monoamine-releasing agent, is a prime candidate in this context. In people with cocaine addiction, damphetamine in long-acting, sustained-release formulations is both safe and effective in reducing cocaine use (Grabowski et al., 2001, 2003; Greenwald et al., 2010; Lile et al., 2020; Nuijten et al., 2016; Rush et al., 2010). D-amphetamine also reduces cocaine use in nonhuman primates (Czoty et al., 2010, 2011; Lile et al., 2020; Negus, 2003; Negus & Mello, 2003) and laboratory rats (Abreu, 1988; Allain et al., 2021; Chiodo et al., 2008; Chiodo & Roberts, 2009; Morgane et al., 2013; Siciliano et al., 2018; Zimmer et al., 2014).

Cocaine addiction afflicts both women and men, and studying addiction treatments in both female and male subjects is especially important because the sexes can differ in the transition to addiction (Becker, 2017; Becker & Koob, 2016; Carroll & Lynch, 2016). Compared to men, women are likely to consume more cocaine, can have a shorter latency from first cocaine use to abuse, and can be more vulnerable to craving and relapse after abstinence [(Elman et al., 2001; Haas & Peters, 2000; McCance-Katz et al., 1999; McKay et al., 1996) but see (Nicolas et al., 2021)]. Similar sex differences are also reported in rats (Algallal et al., 2020; Becker & Koob, 2016; Calipari et al., 2017; Kawa & Robinson, 2018; Lynch & Carroll, 2000; Lynch, 2017; Nicolas et al., 2021; Roberts et al., 1989; Roth & Carroll, 2004). Some, but not all, of these sex differences involve estradiol. For instance, it has been suggested that estradiol might influence the initial acquisition of cocaine-taking behaviour, but once addiction-like behaviour is established, sex hormones are less influential (Becker & Koob, 2016; Perry et al., 2015). Thus, sex differences in the response to cocaine might also involve organisational differences in the brain (Becker & Koob, 2016). This further highlights the need to study both females and males.

Still, virtually all rat studies on the anti-addiction effects of d-amphetamine have included males only (Allain et al., 2021; Chiodo et al., 2008; Chiodo & Roberts, 2009; Siciliano et al., 2018; Zimmer et al., 2014). In parallel, some studies in humans and nonhuman primates have included females, but results were not analysed by sex (Grabowski et al., 2001; Greenwald et al., 2010; Lile et al., 2020; Negus & Mello, 2003; Rush et al., 2010, 2009), presumably because many of these studies were not sufficiently powered to do so. Thus, in investigating the effects of d-amphetamine maintenance treatment on cocaine use, studies examining outcome specifically in females are lacking (Prendergast et al., 2014).

To our knowledge, only Siciliano et al. (2019) have published a report on the potentially therapeutic effects of d-amphetamine in female animals self-administering cocaine. They found that an acute d-amphetamine injection reversed the changes in striatal dopamine transporter function produced by cocaine self-administration. This is an important first step in characterizing the effects of d-amphetamine on cocaine-induced plasticity in females. However, the effects of d-amphetamine on actual cocaine use were not examined. Siciliano et al. (2019) also studied the effects of *acute* d-amphetamine. While *chronic* d-amphetamine administration reduces cocaine reinforcement (Allain et al., 2021; Chiodo et al., 2008; Chiodo & Roberts, 2009; Czoty et al., 2010, 2011; Grabowski et al., 2001, 2003; Greenwald et al., 2010; Lile et al., 2020; Negus, 2003; Negus & Mello, 2003; Nuijten et al., 2016; Rush et al., 2010; Shearer et al., 2003; Siciliano et al., 2018), acute d-amphetamine administration can *increase* cocaine taking and seeking (Barrett et al., 2004; Gerber & Stretch, 1975).

Thus, here we sought to determine the effects of chronic d-amphetamine treatment on cocaine-taking and -seeking behaviour in female rats. To this end, in both previously cocaine-naïve and cocaine-experienced rats, we assessed the effects of damphetamine maintenance treatment on cocaine intake, cocaine-induced psychomotor activity, responding for cocaine under a progressive ratio (PR) schedule, cocaine seeking under extinction and drug-primed reinstatement of extinguished cocaine-seeking behaviour. We evaluated these effects after discontinuing d-amphetamine maintenance treatment because in male rats, d-amphetamine can reduce cocaine use long after treatment cessation (Allain et al., 2021; Chiodo et al., 2008; Chiodo & Roberts, 2009; Siciliano et al., 2018). This suggests therapeutic efficacy that might not require continuous d-amphetamine exposure.

## 2. Materials & Methods

### 2.1 Animals

The animal care committee of the Université de Montréal approved all experimental procedures, and these followed the guidelines of the Canadian Council on Animal Care. A total of 84 adult female Wistar rats (150-175 g upon arrival, Charles River Laboratories, Saint Constant, Quebec, Canada) were housed 1/cage, under a reverse dark/light cycle (12 h/12 h and light off at 8 h30 AM). Self-administration training and testing occurred during the dark phase of the rats’ circadian cycle. Rats had free access to water. During the first three days after arrival, rats had free access to food. From the fourth day on, food was restricted (20 g/day throughout, except during food training where rats received 13 g/day and on the day before and after catheter implantation surgery, where rats received 30 g/day). Giving 20 g/day corresponds to 81% of what adlibitum fed female rats of similar age eat, and this food-restriction paradigm is mild enough to enable rats to gain weight over time (Algallal et al., 2020). Compared to *ad libitum* feeding, mild food restriction produces healthier rats (Martin et al., 2010). In drug self-administration studies, such mild food restriction is often used to facilitate the acquisition of initial operant responding for food reward and to reduce the total quantity of cocaine used (Ahmed & Koob, 1999; Algallal et al., 2020; Robinson et al., 2001).

### 2.2 Apparatus and drugs

We used standard operant cages (Med Associates, St Albans, VT) housed in a sound-attenuating cabinet and equipped for both food pellet and intravenous drug delivery. Each cage contained two retractable levers. Pressing one lever (active) produced food or intravenous cocaine and pressing the other (inactive) lever had no programmed consequences. At the beginning of each session, levers were inserted into the cage and the house light was turned on. During each reward delivery and ensuing 20-s time-out period, the light above the active lever was illuminated and both levers were retracted. At the end of the timeout period, the lever light was turned off, and both levers were inserted back into the cage. Each cage also contained four horizontally aligned infrared photocell beams to measure locomotor behaviour during self-administration sessions. Locomotion was recorded as photocell beam breaks/min. Cocaine hydrochloride (COC; Medisca Pharmaceutique, St-Laurent and Galenova, St-Hyacinthe, Quebec, Canada) was dissolved in 0.9% saline and filtered with Corning bottle-top filters (0.22 μm PES membrane; Fisher Scientific, Whitby, ON, Canada). D-amphetamine (Sigma-Aldrich, Dorset, UK) was dissolved in 0.9% saline. An osmotic minipump (Alzet model 2ML2; Durect Corporation, Cupertino, California) prepared to deliver 5 mg/kg/day d-amphetamine for 14 days was implanted subcutaneously in each d-amphetamine-treated rat. This d-amphetamine treatment regimen significantly decreases cocaine-seeking and -taking behaviours, in male rats (Allain et al., 2021; Chiodo et al., 2008; Siciliano et al., 2019; Zimmer et al., 2014). To prepare a 5 mg/kg/d d-amphetamine solution for the 14-day treatment period, rats were weighed before minipump implantation and we then estimated body weight mid-way through the projected 14-day d-amphetamine treatment (i.e., day 7 of treatment). Actual body weights on day 7 of d-amphetamine treatment indicated that rats were receiving 4.85 mg/kg/day d-amphetamine.

### 2.3 Food self-administration training

In daily 1-h sessions, rats were trained to lever press for food pellets (45 mg, banana-flavoured, grain-based pellets and F0165 bioServ grain pellets VWR, Town of Mount-Royal, Quebec, Canada). Responding was first reinforced under a fixed ratio 1 schedule (FR1). A 20-s timeout period followed each food delivery. Once rats earned at least 20 pellets/session during 2 sessions, they proceeded to a FR3 schedule, until they earned at least 20 pellets/session on 2 sessions. All rats acquired food self-administration behaviour.

### 2.4 Intravenous catheter implantation

After food self-administration training, rats were implanted with a catheter into the right jugular vein (Samaha et al., 2011; Weeks, 1962). Rats were anesthetised with isoflurane (5 % for induction and 2 % for maintenance) and received a perioperative intramuscular injection of penicillin (0.02 ml Derapen, 300mg/ml; CDMV, Saint-Hyacinthe, QC, Canada) and a subcutaneous injection of the analgesic Carprofen Rimadyl (0.03 ml of a 50 mg/ml solution; CDMV, QC, Canada). Rats were left to recuperate for 14-17 days before further behavioural testing. During recovery, incision sites were cleaned with chlorexidine daily. Starting from the day after catheter implantation, catheters were flushed daily with 0.05 ml saline, or 0.1 ml of a 0.2 mg/ml heparin/saline solution mixed with 2 mg/ml of baytril (CDMV, St Hyacinthe, Quebec, Canada) on alternate days.

### 2.5 Cocaine self-administration training

Following recovery from catheter implantation, rats were trained to self-administer i.v. cocaine (0.25 mg/kg/infusion, delivered over 5 s) under FR3, during 1-h daily sessions. During each session, cocaine was continuously available. During each 5-s cocaine infusion, the levers retracted and the light above the active lever was turned on. We considered that rats acquired reliable cocaine self-administration when, for two consecutive sessions, they took at least 6 injections/session and pressed twice more on the active versus inactive lever.

### 2.6 Subcutaneous minipump implantation

Under isoflurane anesthesia (5% for induction and 2% for maintenance), a d-amphetamine-filled minipump was implanted subcutaneously in ‘COC + A’ rats (Exp. 1) and in ‘COC/COC + A’ rats (Exp. 2). ‘COC’ rats (Exp. 1) and ‘COC/COC’ rats (Exp. 2) received a sham surgery consisting of an incision and sutures. We did not use saline-filled minipumps in control rats, because in males, sham-operated rats and rats with a saline-filled minipump show similar cocaine self-administration behaviour (Allain et al., 2021).

### 2.7 Intermittent access (IntA) cocaine self-administration

Rats received 14 daily sessions (5 h/session) during which they could self-administer cocaine under intermittent-access conditions (Zimmer et al., 2012). During each daily session, cocaine (0.25 mg/kg/infusion, delivered over 5 s) was available in 5-min periods, separated by 25-min periods during which levers were retracted and cocaine was unavailable. Each 5-h session thus consisted of 10 cocaine ON/OFF cycles. This intermittent-access procedure is especially effective in producing addiction-relevant patterns of cocaine use, and it also models the intermittency of human cocaine (Allain et al., 2015; Kawa et al., 2019b; Samaha et al., 2021).

### 2.8 Progressive ratio

We assessed responding for cocaine under a progressive ratio (PR; Response ratio = [5e ^(*injection number x* 0.2)^)] - 5) schedule of reinforcement (Richardson & Roberts, 1996). The rats responded for 3 cocaine doses (0.063, 0.125 and 0.25 mg/kg/infusion; 1 dose/ session, in counterbalanced order). PR sessions ended after 5 hours or when 1 h elapsed since the last self-administered injection.

### 2.9 Extinction

All rats received 10 daily extinction sessions (2 h/session). During these sessions, the rats were tethered to the infusion apparatus, and pressing the active lever produced the discrete environmental cues previously associated with cocaine (illumination of the lever above the active lever, retraction of both levers and activation of syringe pumps), under FR3. However, no cocaine was available.

### 2.10 Cocaine-induced reinstatement

Sessions were like extinction sessions, except that rats now received i.p. injections of saline (session 1) and cocaine (7.5 mg/kg for session 2) right before being placed into the operant cages for 2 h. During these sessions, the houselight in each cage was off and the rats were not tethered to the infusion system.

### 2.11 Experiment 1: Effects of d-amphetamine treatment in previously cocaine-naïve rats

Fig. 1A shows the sequence of experimental events. After self-administration training, rats were assigned to the COC (n = 20) or COC + A (n = 17) groups, such that average numbers of infusions, and active and inactive lever presses during the last two days of self-administration training were held constant across groups. COC + A rats were then implanted with a d-amphetamine-containing minipump, and COC rats received a sham surgery. One day later, all rats received the first of 14 IntA cocaine sessions (1 session/day). The day after the last session, minipumps were removed from COC + A rats, and COC rats received a second sham surgery, so that both groups received the same number of surgeries. Two days later, we measured responding for cocaine under a PR schedule of reinforcement (0.063, 0.125 and 0.25 mg/kg/infusion, 1 session/dose counterbalanced). After PR testing, all rats were given 10 extinction sessions (1 session/day). Finally, we measured cocaine-induced (0 and 7.5 mg/kg, i.p) reinstatement of drug-seeking behaviour.

**Figure 1.**
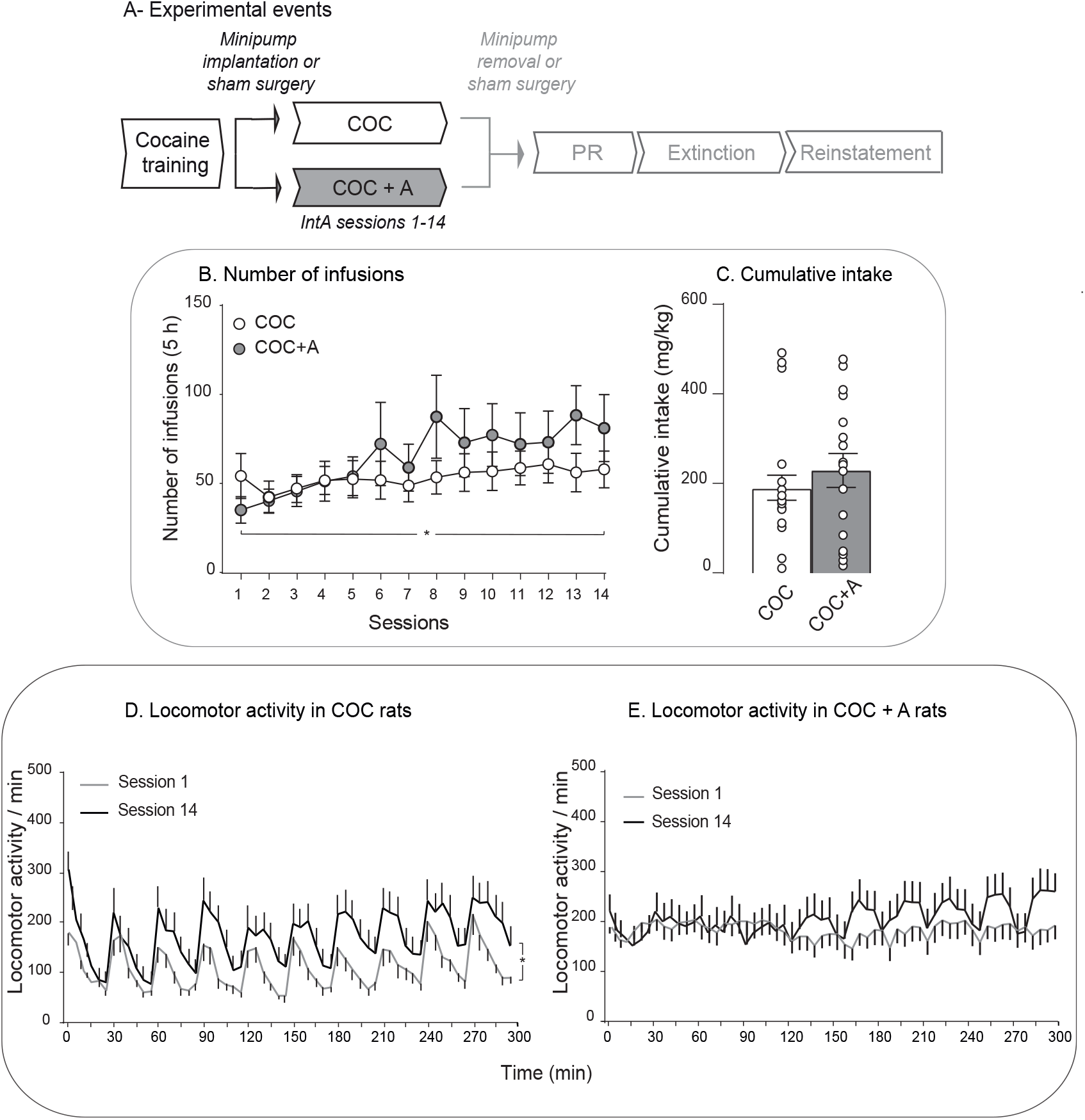
D-amphetamine treatment did not change ongoing cocaine intake but it prevented both spikes in locomotor activity during cocaine self-administration sessions and the development of cocaine-induced psychomotor sensitization. (A) The sequence of experimental events for experiment 1. After implantation of an intravenous catheter into the jugular vein, female rats were trained to self-administer intravenous cocaine (0.25 mg/kg/infusion, injected over 5 s) during daily, 1-h sessions. Rats then self-administered cocaine during 14, 5-h intermittent access sessions. Some rats received concomitant d-amphetamine treatment (COC + A rats; 5 mg/kg/day, via s.c. minipump). During d-amphetamine treatment/intermittent cocaine self-administration, we measured cocaine intake and locomotion. After discontinuation of d-amphetamine treatment/intermittent cocaine self-administration, we measured responding for cocaine under a progressive ratio schedule (PR), responding under extinction, and cocaine-primed reinstatement of extinguished cocaine-seeking behaviour. (B) COC + A rats and COC rats took similar amounts of cocaine across the 14 self-administration sessions, and both groups also escalated their intake over sessions. (C) Cumulative intake was comparable between the groups. (D) COC rats increased their horizontal locomotor activity over sessions, suggesting that COC rats developed psychomotor sensitization to self-administered cocaine. Intermittent cocaine intake also produced spikes in locomotion across the 5-h sessions. (E) COC + A showed no significant increase in locomotion across sessions, suggesting that d-amphetamine suppressed the development of psychomotor sensitization. D-amphetamine also suppressed the spikes in locomotion normally produced by intermittent cocaine self-administration. Data are mean ± SEM. n = 20 for the COC group and n = 17 for the COC + A group. In (B), * p < 0.05, Session effect. In (D), * p < 0.05, Session effect and Session x Time interaction effect.

### 2.12 Experiment 2: Effects of d-amphetamine treatment in cocaine-experienced rats

In Experiment 1 we assessed the effects of d-amphetamine treatment in previously cocaine-naïve rats, as this allows for comparison with previous studies, conducted in males (Allain et al., 2021; Chiodo et al., 2008; Chiodo & Roberts, 2009; Siciliano et al., 2018). In Experiment 2, we assessed d-amphetamine’s effects in rats with a history of cocaine intake, because in the clinic, individuals presenting for cocaine addiction treatment will be cocaine experienced. Fig. 4A shows the sequence of experimental events. After self-administration training, all rats (n = 26) self-administered cocaine during 14 IntA sessions (Sessions 1-14). Three days later, we established baseline levels of responding for cocaine under a PR schedule of reinforcement (0.063, 0.125 and 0.25 mg/kg/infusion, 1 session/dose counterbalanced). After the last PR session, rats were assigned to a ‘COC/COC’ (n = 7) or ‘COC/COC + A’ group (n = 9), such that average number of infusions taken over the 14 intermittent sessions, ratios reached for cocaine (0.063, 0.125 and 0.25 mg/kg/infusion), active lever presses and number of infusions earned during PR sessions were held constant across groups. COC/COC + A rats received a d-amphetamine-filled minipump. COC/COC rats were sham-operated. The day after this surgery, all rats received 14 additional IntA sessions (Sessions 15-28). The day after session 28, minipumps were removed from COC/COC + A rats and COC/COC rats were sham-operated once more. Two days after, we measured responding for cocaine (0.063, 0.125 and 0.25 mg/kg/infusion, 1 session/dose, counterbalanced) under PR. The day after the last PR session, rats received ten, 2-h extinction sessions, followed by cocaine-induced (0 and 7.5 mg/kg, i.p) reinstatement tests.

### 2.13 Catheter patency

In each experiment, we verified catheter patency after the last IntA cocaine session and again after the last PR session, by administering an i.v. infusion of propofol (10 mg/mL; 0.1 mL/rat; CDMV, St-Hyacinthe, QC). Rats that became ataxic within 10 seconds after infusion were deemed to have a functional catheter, and their data were included in the results. Data from rats that did not become ataxic (31 rats across experiments 1 & 2) were excluded from analysis. This attrition rate is greater than what our laboratory typically sees in intravenous drug self-administration studies in male rats. There can be a greater incidence of i.v. cathether-related malfunctions in female vs male rats (Allain et al., 2022).

### 2.14 Statistical analysis

Data were analysed with GraphPad Prism 8 and the alpha level was set at *p* ≤ 0.05. During IntA, two-way analysis of variance (ANOVA) was used to analyse group differences in the number of self-administered cocaine infusions (Group x Session; the latter as a within-subjects variable). Group differences in cumulative intake after IntA sessions were analysed using unpaired t-tests. Two-way, repeated measures ANOVA was used to analyse locomotor activity across IntA sessions (for within-groups comparisons; Session x Time in min; both as within-subjects variables; for between-groups comparisons; Group x Time in min; the latter as a within-subjects variable). Two-way ANOVA was also used to analyse group differences in lever pressing during PR sessions (Group x Dose; the latter as a within-subjects variable). To analyse the relationship between cumulative cocaine intake during IntA sessions and number of infusions earned during PR testing, we assessed goodness-of-fit (r^2^) of the linear regression between the two measures. We analysed group differences in lever pressing during extinction and reinstatement tests with three-way ANOVA (for extinction: Group x Lever x Session; the latter two as within-subjects variables. For reinstatement: Group x Lever x Cocaine dose; the latter two as within-subjects variables). Data in Fig. are expressed as mean values ± S.E.M.

## 3. Results

### 3.1 Experiment 1: Effects of d-amphetamine treatment in previously cocaine-naïve rats

Figure 1B shows that across COC and COC + A groups, rats escalated their cocaine intake over the IntA sessions (Session effect, F (3.7, 129.5) = 2.637, *p* = 0.04), with no group differences in intake across sessions (all *P*’s > 0.05). Cumulative cocaine intake after the 14 self-administration sessions was also similar across groups (Fig. 1C; *p* > 0.05). Thus, d-amphetamine treatment did not significantly change ongoing cocaine intake. Fig. 1D shows that in COC rats, cocaine-induced locomotion increased from Session 1 to 14 (Session effect, F (1, 19) = 5.84, *p* = 0.03; Session x time interaction effect, F (59, 1121) = 1.367, *p* = 0.04). This suggests that COC rats developed psychomotor sensitization to cocaine. In contrast COC + A rats did not show a significant change in locomotion over sessions (Fig. 1E; all *P*’s > 0.05). Thus, to the extent that an increase in locomotor activity across sessions reflects the development of psychomotor sensitization, these findings suggest that d-amphetamine prevented the development of psychomotor sensitization otherwise produced by IntA cocaine intake. Of note, psychomotor sensitization can also involve an increased incidence of stereotypic behaviours with repeated drug treatment, and we did not measure stereotypy here. On Session 1, COC + A rats showed greater locomotion than COC rats did (gray curve in Fig. 1D vs. that in 1E; Group effect, F (1, 35) = 6.91, *p* = 0.01). However, the two groups showed similar levels of locomotion by Session 14 (black curve in Fig. 1D vs. that in 1E; *p* > 0.05). Thus, initially, d-amphetamine maintenance treatment during cocaine self-administration can increase psychomotor activity relative to cocaine self-administration alone, but over time, psychomotor activity becomes similar under the two conditions. There were also *qualitative* differences in cocaine-induced locomotion between the groups. During Sessions 1 and 14, COC rats showed a spiking pattern of locomotion, whereas COC + A rats showed comparatively flattened locomotion curves (Session 1, gray curve in Fig. 1D vs. that in 1E; Time effect, F (8.203, 287.1) = 3.03; Time x Group effect F (59, 2065) = 3,84; All *P*’s < 0.0001; Session 14, black curve in 1D vs. that in 1E; F (4.759, 166.6) = 3.72; *p* = 0.004; Time x Group effect F (59, 2065) = 3,32; *p* < 0.0001). Thus, d-amphetamine treatment attenuated the spikes in locomotor activity otherwise produced by IntA cocaine intake. In summary, the findings suggest that d-amphetamine both prevented psychomotor sensitization to self-administered cocaine and changed the temporal kinetics of cocaine induced locomotion [see also (Allain et al., 2021)].

#### 3.1.1 Responding for cocaine under a PR schedule of reinforcement

D-amphetamine treatment was discontinued after the 14 IntA sessions, and 3 days later, we assessed responding for cocaine under a PR schedule (Fig. 2B). Across groups, rats lever pressed more for larger cocaine doses (Dose effect, F (1.885, 65.97) = 13.84, *p* < 0.0001). There were also group differences in responding for cocaine. Across doses, COC + A rats lever pressed less for cocaine than COC rats did (Group effect, F (1, 35) = 4.18, *p* = 0.048, Fig. 2B). Thus, d-amphetamine treatment significantly reduced the later motivation to obtain cocaine. Figs. 2C-E show that in both groups, higher levels of cumulative cocaine intake across the previous 14 IntA sessions predicted greater responding for the drug during PR tests (COC rats; 0.063 mg/kg/infusion, r^2^ = 0.33; 0.125 mg/kg/infusion, r^2^ = 0.33; 0.25 mg/kg/infusion, r^2^ = 0.17; COC + A rats; 0.063 mg/kg/infusion, r^2^ = 0.63; 0.125 mg/kg/infusion, r^2^ = 0.60; All *P*’s ≤ 0.008; except for COC rats at 0.25 mg/kg/infusion, *p* > 0.05). Thus, rats that self-administered more cocaine in the past also subsequently worked harder to obtain cocaine under a PR schedule of reinforcement.

**Figure 2.**
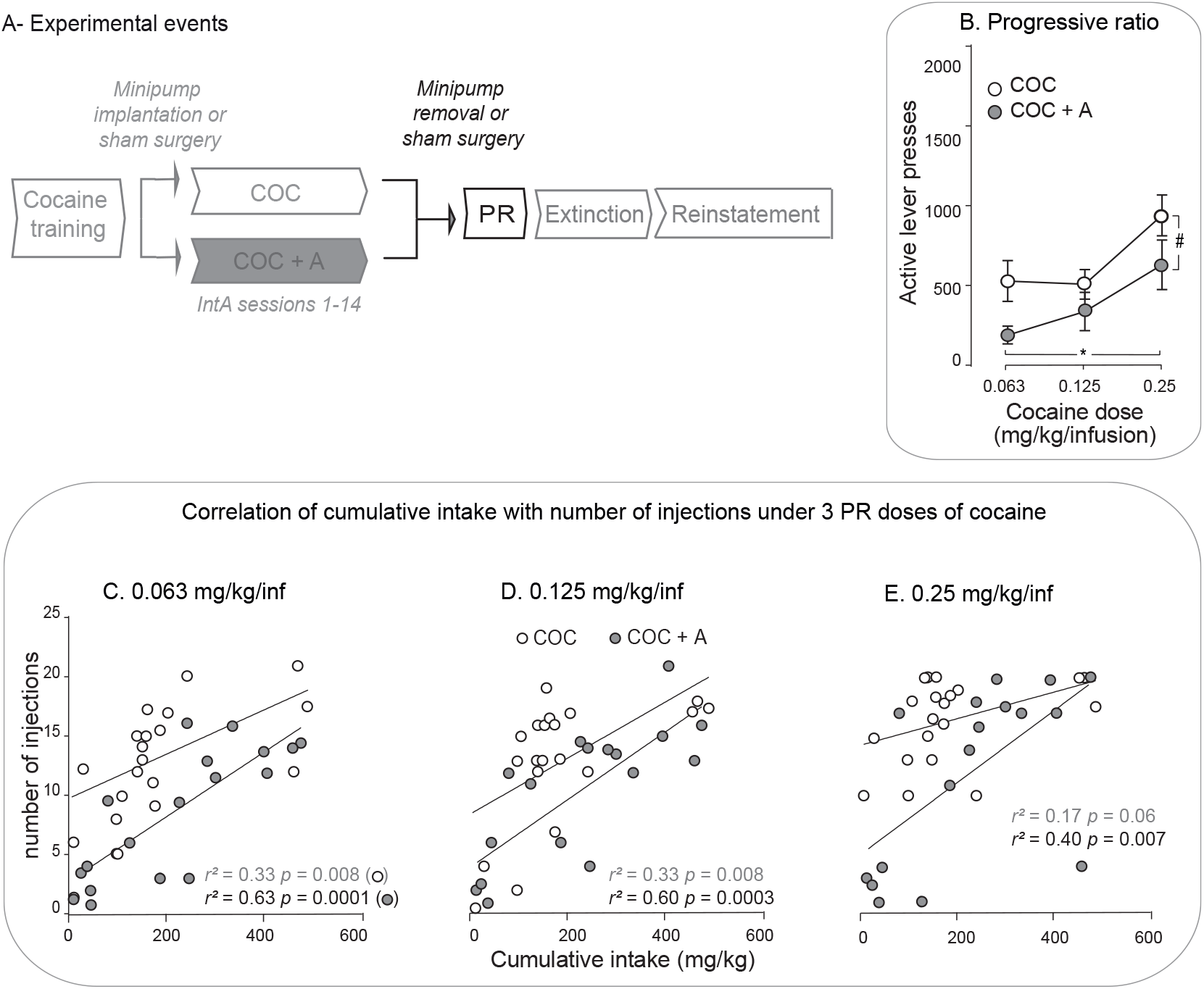
D-amphetamine treatment during intermittent cocaine self-administration decreased later responding for cocaine under a progressive ratio schedule of reinforcement and cumulative cocaine intake during intermittent self-administration sessions was positively correlated with cocaine infusions earned under progressive ratio. (A) Sequence of events for experiment 1. (B) During progressive ratio sessions, responding was dose-dependent, and COC + A rats responded less for cocaine than COC rats did. This indicates that d-amphetamine treatment reduced later responding for cocaine under progressive ratio. (C-E) Correlations between average cumulative cocaine intake during the 14 intermittent access sessions and average number of cocaine infusions earned during progressive ratio sessions at (C) 0.063 mg/kg/infusion, (D) 0.125 mg/kg/infusion and (E) 0.25 mg/kg/infusion cocaine. In both COC + A and COC rats, more cumulative cocaine intake predicted more responding for cocaine at all doses tested under a progressive ratio schedule, save for the 0.25 mg/kg dose in the COC rats, where the correlation was nonsignificant. Data are mean ± SEM. n = 20 for the COC group and n = 17 for the COC + A group. In (B), # p < 0.05, Group effect; * p < 0.0001, Dose effect.

#### 3.1.2 Extinction and reinstatement

After PR testing, rats received 10 extinction sessions, during which pressing the active lever produced the discrete cues previously associated with cocaine, but not cocaine itself. Fig. 3B shows that during these sessions, both COC + A and COC rats pressed more on the active versus inactive lever (Lever effect, F (1, 35) = 51.5, *p* < 0.0001), and both groups also progressively reduced their pressing behaviour on the active lever across sessions (Session effect, F (9, 315) = 13.36; Lever x Session, F (9, 315) = 9.87; All *P*’s < 0.0001). There were also group differences in responding. COC + A rats responded less than COC rats did under extinction (Lever x Session x Group, F (9, 315) = 2.91, *p* = 0.04; Group effect, F (1, 35) = 6.22, *p* = 0.02; Fig. 2B). Thus, receiving d-amphetamine maintenance treatment decreased later cocaine-seeking behaviour. Because d-amphetamine treatment was stopped 9 days prior to the first extinction session, the findings show that d-amphetamine treatment produced a long-lasting decrease in the incentive value of cocaine, of the cocaine-paired cues/context or both.

**Figure 3.**
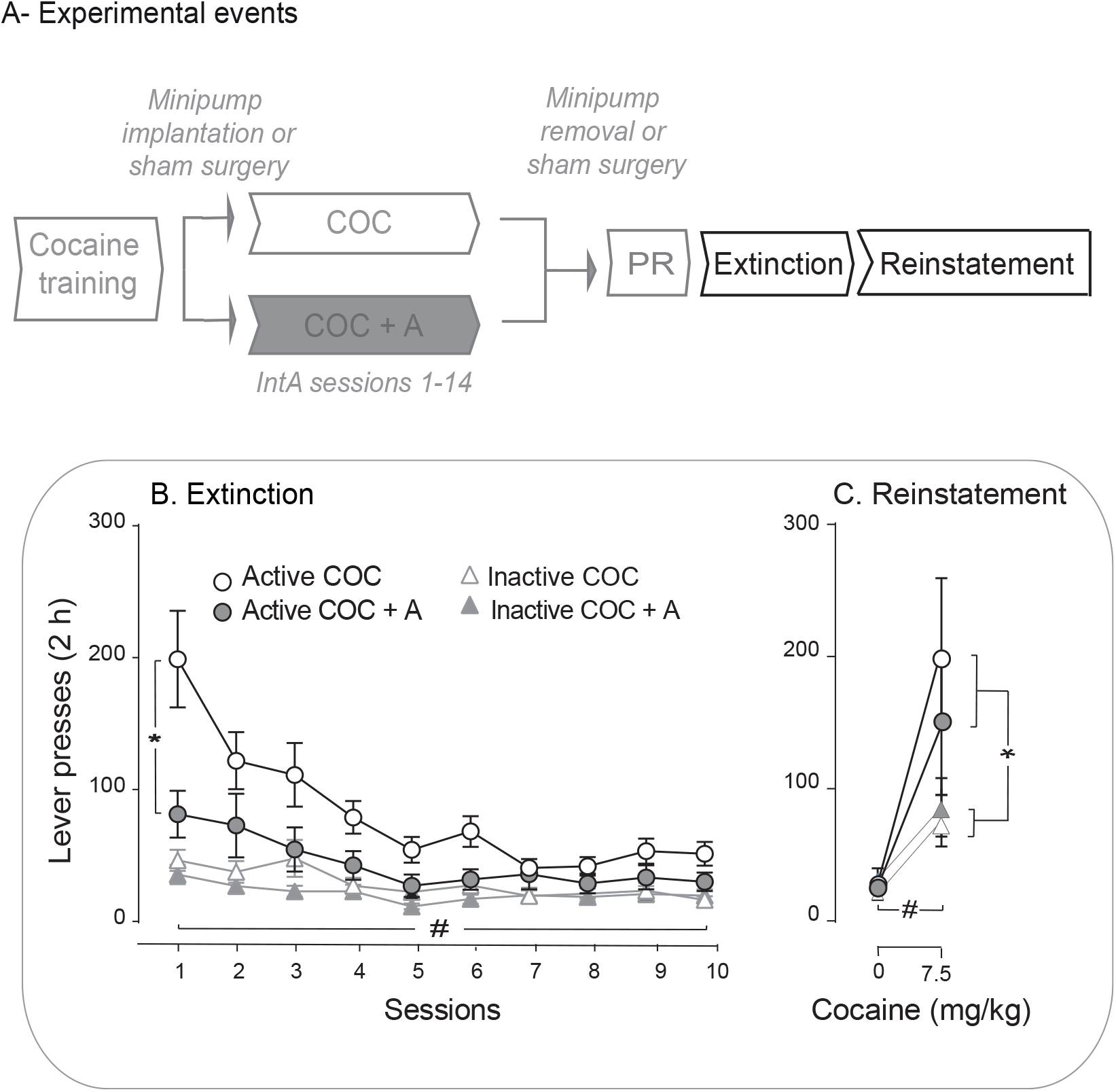
D-amphetamine treatment during intermittent cocaine self-administration decreased later cocaine seeking. (A) Sequence of events for experiment 1. (B) Under extinction conditions, both groups extinguished responding over sessions, and COC + A rats showed less cocaine-seeking behaviour compared to COC rats. Thus, d-amphetamine treatment suppressed later cocaine seeking under extinction conditions. (C) Both groups showed cocaine-induced reinstatement of extinguished cocaine seeking, and there were no group differences in responding. Data are mean ± SEM. n = 20 for the COC group and n = 17 for the COC + A group. In (B), * p < 0.05, Group effect; # p < 0.0001, Session effect. In (C), * p < 0.05, Lever effect, p < 0.0001 Dose effect.

**Figure 4.**
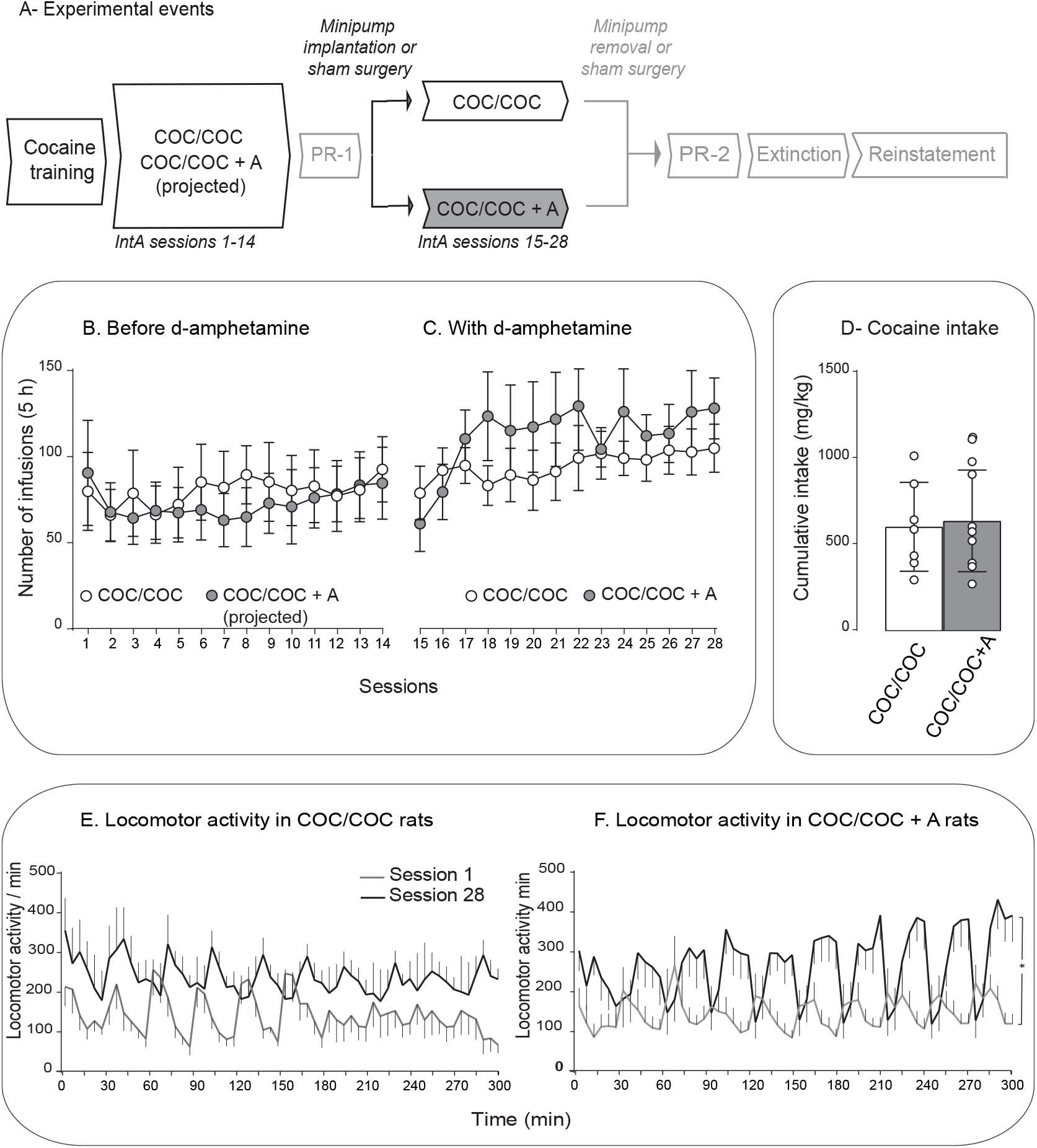
bfD-amphetamine treatment did not influence cocaine intake in already cocaine-experienced rats.(A) Sequence of events in experiment 2. After implantation of an intravenous catheter into the jugular vein, female rats were trained to self-administer intravenous cocaine (0.25 mg/kg/infusion, injected over 5 s) during daily, 1-h sessions. Rats then self-administered cocaine during 14, 5-h intermittent access sessions (Sessions 1-14). We then established baseline levels of responding for cocaine under a progressive ratio schedule of reinforcement (PR-1). The rats then received 14 more intermittent access sessions (Sessions 15-28). Some rats received concomitant d-amphetamine treatment during these additional cocaine self-administration sessions (COC/COC + A rats; 5 mg/kg/day, via s.c. minipump). During all intermittent cocaine self-administration sessions, we measured cocaine intake and locomotion. After the 28th self-administration session, we discontinued d-amphetamine treatment, and we measured responding for cocaine under a progressive ratio schedule once again (PR-2). Finally, we also measured responding under extinction, and cocaine-primed reinstatement of extinguished cocaine-seeking behaviour. (B-C) COC rats and COC/COC + A rats self-administered a similar number of cocaine injections across the 28 intermittent access sessions. (D) Cumulative cocaine intake across the 28 intermittent access sessions was similar between the groups. (E) COC/COC rats showed no significant change in locomotion from session 1 to session 28. (F) COC/COC + A rats showed significantly more locomotion on Session 28 vs. Session 1. Data are mean ± SEM. n = 7 for the COC/COC group and n = 9 for the COC/COC + A group. In (F), * p < 0.05, Session effect, p < 0.0001, Session x Time interaction effect.

After extinction sessions, rats were tested for cocaine-primed reinstatement of drug-seeking behaviour (Fig. 3C). Across groups, cocaine increased lever-pressing behaviour, and did so preferentially on the previously cocaine-associated, active lever (Dose effect, F (1, 35) = 25.05, *p* < 0.0001; Lever effect, F (1, 35) = 5.59, *p* = 0.02; Lever x Dose, F (1, 35) = 6.39, *p* = 0.02). There were no group differences in cocaine-induced reinstatement of drug-seeking behaviour. Thus, prior d-amphetamine treatment did not change later cocaine-primed relapse behaviour.

### 3.2 Experiment 2: Effects of d-amphetamine treatment in previously cocaine-experienced rats

As shown in Fig. 4A, a new group of rats received 14 initial IntA sessions (sessions 1-14), without d-amphetamine, followed by baseline PR tests (i.e., PR-1). All rats then received 14 more IntA sessions (Sessions 15-28). Some rats received d-amphetamine treatment (COC/COC + A rats) during these additional 14 sessions, and other rats did not (COC/COC rats). After Session 28, d-amphetamine treatment was ceased and all rats were tested once more under a PR schedule of reinforcement (PR-2), followed by extinction and cocaine-primed reinstatement tests.

#### 3.2.1 IntA cocaine self-administration

This cohort of rats did not significantly escalate their cocaine intake across IntA sessions (Figs. 4B and 4C; All *P*’s > 0. 05). During sessions 1-14 (Fig. 4B), and again during sessions 15-28 (Fig. 4C), rats in the two groups also took similar amounts of cocaine (All *P*’s > 0. 05). Cumulative cocaine intake across the 28 sessions was therefore similar across the groups (Fig. 4D; *p* > 0. 05). Thus, as in Experiment 1, d-amphetamine treatment did not significantly influence ongoing cocaine intake. To determine whether rats developed psychomotor sensitization across the 28 IntA self-administration sessions, we compared locomotion on IntA session 1 vs. 28 in each group. In each group, there was a significant Time (min) x Session interaction effect when comparing locomotion on IntA session 1 vs. 28 (Time (min) x Session interaction; Fig. 4E; F (59, 708) = 1.78, *p* = 0.0005; Fig. 4F; F (59, 944) = 6.5, *p* < 0.0001). This suggests an increased psychomotor response to cocaine over sessions. COC and COC+A rats showed comparable levels of locomotion on Session 1, when both groups were taking cocaine without d-amphetamine co-treatment (F (1, 14) = 0.004, *p* = 0.95), and on Session 28, when COC+ A rats were now on d-amphetamine (F (1, 14) = 0.15, *p* = 0.71). This suggests that both groups developed similar levels of psychomotor sensitization across the 28 IntA sessions. However, there was a significant Time (min) by Group interaction effect on Session 28 (black curve in Fig. 4E vs. that in Fig. 4F; F (59, 826) = 2.40, *p* < 0.0001). This suggests that d-amphetamine maintenance therapy changed the temporal kinetics of the locomotor response to cocaine. To examine this further, we compared the groups on locomotor activity during the 5-min cocaine ON periods and during the 25-min cocaine OFF periods on Session 28. D-amphetamine decreased locomotion during the 5-min cocaine ON periods (Figure S1; p < 0.0001) and it increased locomotion during the 25-min cocaine OFF periods (Figure S1; p < 0.0001). The decrease in locomotion during the cocaine ON periods could reflect stereotypy, as locomotor activity and stereotypy can be mutually exclusive behaviours. However, we did not measure stereotypy here. Whatever the case may be, the results suggest that in rats with a prior history of cocaine intake (Sessions 1-14), d-amphetamine treatment did not prevent the development of psychomotor sensitization to self-administered cocaine.

#### 3.2.2 Responding for cocaine under a PR schedule

We assessed responding for cocaine under a PR schedule of reinforcement after IntA sessions 1-14 (‘PR1’ in Figs. 5B-I), and again after IntA sessions 15-28 (‘PR2’). During sessions 15-28, COC/COC +A rats received concomitant d-amphetamine treatment. COC/COC rats remained d-amphetamine-free. Figs. 5B-I show responding for cocaine during these PR tests. COC/COC rats responded more for larger cocaine doses (Fig. 5B, Dose effect, F (1.196, 14.35) = 10.26, *p* = 0.005), and they showed similar responding for cocaine during PR-1 and PR-2 tests (All *P*’s > 0.05; see individual data points in Figs. 5C-E). COC/COC + A rats also lever pressed more for larger cocaine doses (Fig. 5F; Dose effect, F (2,48) = 9.35, *p* = 0.0004). Importantly, prior d-amphetamine treatment reduced responding for cocaine under a PR schedule, as assessed both within and between subjects. First, COC/COC + A rats responded significantly less for cocaine after d-amphetamine treatment vs. before (Fig. 5F; PR1 vs. PR2, F (1, 16) = 4.72, *p* = 0.04). Second, before d-amphetamine treatment, responding for cocaine was similar between COC/COC and COC/COC + A groups (‘PR-1’ in Fig. 5B vs that in Fig. 5F; All *P*’s > 0.05). However, after receiving d-amphetamine treatment, COC/COC + A rats responded less than COC/COC rats did (‘PR-2’ in Fig. 5F vs that in Fig. 5B; Group effect, F (1, 14) = 6.214, *p* = 0.03). In summary, in rats with a prior history of cocaine intake, d-amphetamine maintenance treatment significantly reduced the later motivation to obtain cocaine.

**Figure 5.**
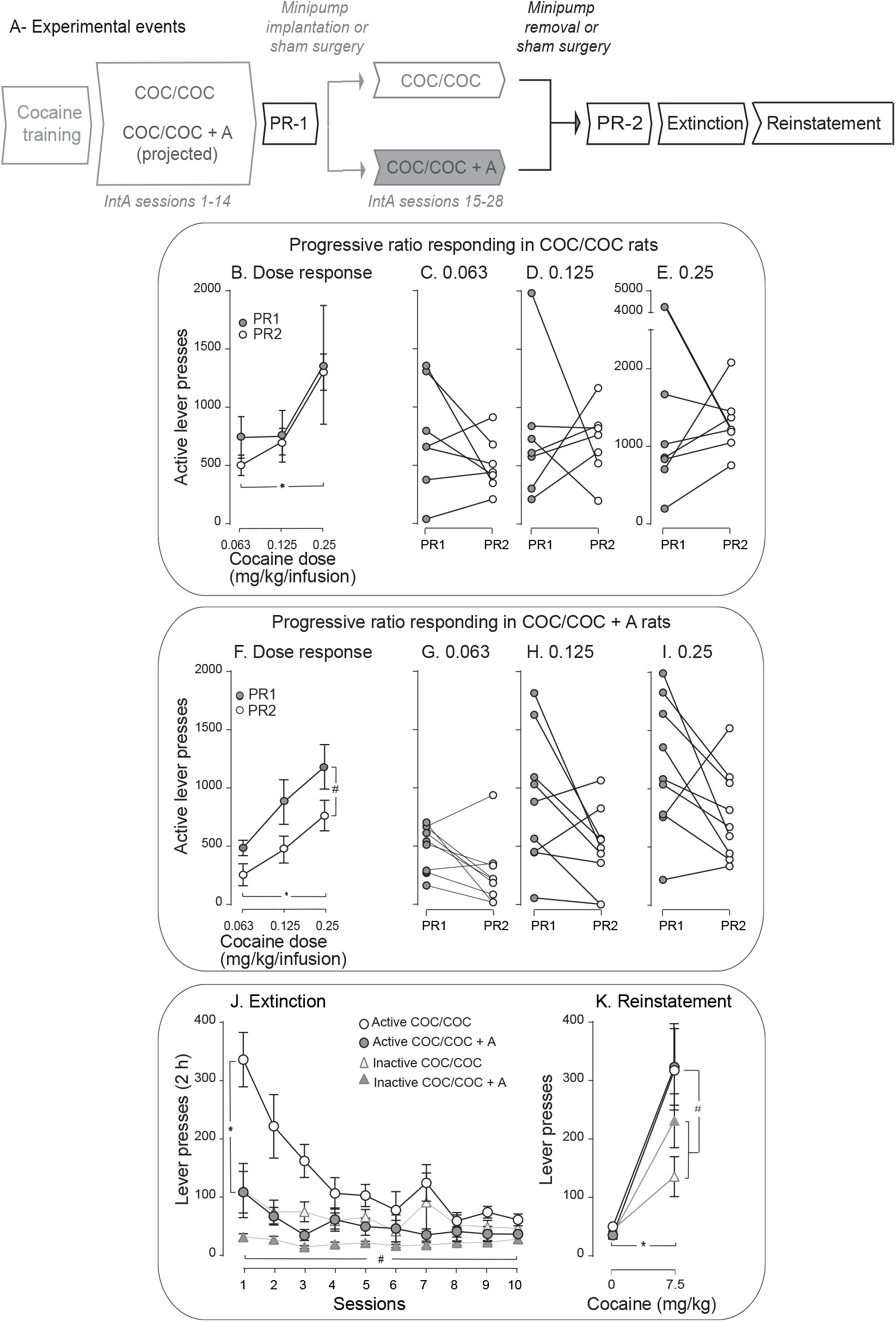
In cocaine-experienced rats, D-amphetamine treatment during intermittent cocaine self-administration sessions decreased later responding for cocaine both under a progressive ratio schedule of reinforcement and under extinction conditions. (A) Sequence of experimental events for experiment 2. (B-E), During progressive ratio sessions, COC/COC rats showed a dose-dependent increase in active lever pressing for cocaine and responding was similar during the PR-1 vs. PR-2 tests. (C-E) Responding for cocaine under progressive ratio in individual COC/COC rats during PR-1 and PR-2 tests. (F-I) COC/COC + A rats showed a dose-dependent increase in active lever pressing for cocaine, and they responded significantly less during PR-2 vs. PR-1 tests. Thus, d-amphetamine treatment decreased later responding for cocaine under a progressive ratio schedule of reinforcement. (G-I), Responding for cocaine under progressive ratio in individual COC/COC +A rats during PR-1 and PR-2 tests. (J) Under extinction conditions, both groups extinguished responding over sessions, and COC/COC + A rats showed less cocaine-seeking behaviour compared to COC/COC rats. Thus, d-amphetamine treatment decreased later cocaine-seeking behaviour under extinction conditions. (K) Both groups showed cocaine-induced reinstatement of extinguished cocaine seeking, and there were no group differences. Data are mean ± SEM. n = 7 for the COC/COC group and n = 9 for the COC/COC + A group. In (B), * p = 0.005, Dose effect. In (F), # p < 0.05, Group effect; * p < 0.0001, Dose effect. In (J), * p < 0.05, Group effect, # p < 0.001, Session effect. In (K), # p < 0.0001, Lever effect, * p = 0.05, Dose effect.

#### 3.2.3 Extinction and reinstatement

Following PR testing, all rats received 10 extinction sessions, followed by a cocaine-primed reinstatement test. Fig. 5J shows that across groups, rats pressed more on the previously cocaine-associated (active) lever vs. on the inactive lever (Lever effect, F (1, 14) = 19.12, *p* = 0.001), and the rats also reduced their leverpressing behaviour across sessions (Session effect, F (9, 126) = 11.88, *p* < 0.0001), with a more pronounced reduction in presses on the previously active vs. inactive lever (Lever x Session effect, F (9, 126) = 12.77, *p* < 0.0001). Thus, rats extinguished their cocaine-seeking behaviour across sessions. There were also group differences in extinction responding. COC/COC + A rats lever pressed significantly less than COC/COC rats did, indicating reduced cocaine-seeking behaviour (Lever x Session x Group interaction, F (9,126) = 4.758, *p* = 0.006; Group effect, F (1, 14) = 14.335, *p* = 0.02; Session x Group, F (9, 126) = 5.321, *p* = 0.006). In summary, in rats that had a history of cocaine use before d-amphetamine treatment, d-amphetamine maintenance therapy significantly reduced later cocaine-seeking behaviour.

Fig. 5K shows that across groups, a priming injection of cocaine (7.5 mg/kg, i.p.) reinstated extinguished cocaine-seeking behaviour (Dose effect, F (1, 14) = 28.6, *p* < 0.0001; Lever effect, F (1, 14) = 4.65, *p* = 0.05; Lever x Dose interaction, F (1, 14) = 6.32, *p* = 0.03). However, there were no group differences in this behaviour. Thus, in rats that were cocaine-experienced rats before being treated with d-amphetamine, d-amphetamine maintenance treatment did not change later cocaine-primed relapse.

## 4. Discussion

Studies in humans and laboratory animals show that D-amphetamine administration can reduce cocaine-seeking and -taking behaviours. These studies have generally reported on effects in males, and it is not clear whether similar effects occur in females. Here female rats had intermittent access to cocaine, with or without d-amphetamine treatment, and we examined features relevant to addiction. These included the development of psychomotor sensitization, high levels of incentive motivation for the drug, as measured using PR procedures, resistance to extinction, and cocaine-triggered relapse-like behaviour after abstinence. We report four main findings. First, in both previously cocaine-naïve and cocaine-experienced rats, d-amphetamine treatment did not significantly change ongoing cocaine intake.

Second, in previously cocaine-naïve rats only, d-amphetamine treatment attenuated the development of psychomotor sensitization otherwise produced by IntA cocaine intake. Third, in both previously cocaine-naïve and cocaine-experienced rats, d-amphetamine treatment reduced incentive motivation for cocaine and cocaine seeking under extinction conditions. Finally, d-amphetamine did not influence cocaine-primed reinstatement of extinguished drug-seeking behaviour. Thus, d-amphetamine treatment reduces several behavioural indices of increased incentive motivation for cocaine in female rats. To the degree that the animal model we used is relevant to cocaine addiction in humans, our findings suggest that continuous d-amphetamine treatment could have long-lasting therapeutic effects in treating cocaine addiction.

### 4.1 Intermittent cocaine intake

We used an IntA cocaine self-administration procedure here (Zimmer et al., 2011, 2012). IntA differs from other self-administration procedures often used to model the transition to addiction, such as Long Access. Long Access involves continuous drug availability during each session. In contrast, during IntA, cocaine is available intermittently, producing peaks and troughs in brain concentrations of the drug (Algallal et al., 2020; Allain et al., 2017b; Allain & Samaha, 2019; Allain et al., 2017a; Zimmer et al., 2012). This more closely models human patterns of cocaine use, which are thought to be intermittent, both between and within periods of use (Allain et al., 2015; Beveridge et al., 2012; Cohen & Sas, 1994; Leri et al., 2004). IntA is also uniquely effective in producing addiction-relevant patterns of drug taking and seeking (Allain et al., 2015; Kawa et al., 2019b; Samaha et al., 2021). IntA drug experience produces persistent psychomotor sensitization (Algallal et al., 2020; Allain et al., 2017b; Carr et al., 2020), long-lasting enhancements in the motivation to obtain cocaine (Algallal et al., 2020; Allain & Samaha, 2019; Calipari et al., 2014a; James et al., 2019), increased persistence of drug taking in the face of an aversive consequence (James et al., 2019; Singer et al., 2018), and very robust cue-induced reinstatement of cocaine seeking (Gueye et al., 2019; James et al., 2019; Kawa et al., 2019a, 2016; Nicolas et al., 2019; Singer et al., 2018). IntA cocaine experience can produce quantitatively different effects in females versus males. With IntA cocaine intake, female rats show more robust psychomotor sensitization than male rats do (Algallal et al., 2020; Carr et al., 2020), and females also more readily develop incentive sensitization, as indicated by earlier and greater increases in the motivation for cocaine (Kawa et al., 2019a) and more cocaine seeking when the drug is unavailable (Nicolas et al., 2019). These observations reflect the female ‘telescoping effect’, whereby in people vulnerable to addiction, women can transition more readily from initiation of drug use to addiction than men do [(Becker, 2017), see also (Nicolas et al., 2021)].

### 4.2 D-amphetamine effects on the response to cocaine in female and male rats

Our findings in female rats here are comparable but not identical to those we reported in male rats in recent work (Allain et al., 2021). We compare across studies with caution. With this caveat in mind, using identical experimental procedures as used here, we found that d-amphetamine attenuated the development of psychomotor sensitization to self-administered cocaine and decreased responding for cocaine during both PR and extinction sessions in male rats (Allain et al., 2021), as observed here in females. However, there are also two main differences in the findings. First, in male rats, d-amphetamine reduced both the development and expression of psychomotor sensitization to IntA cocaine self-administration (Allain et al., 2021). In contrast, here in females d-amphetamine attenuated the *development* of psychomotor sensitization to cocaine in previously drug-naïve rats (Fig. 1D-E), but d-amphetamine did not prevent the *expression* of psychomotor sensitization in already cocaine-experienced rats (Fig. 4E-F). In fact, analysis of locomotor activity in the cocaine-experienced rats showed that relative to controls, d-amphetamine-treated rats have decreased locomotor counts during the cocaine ON periods, and increased locomotor counts during the cocaine OFF periods. This could reflect increased stereotypy during the cocaine ON periods in these rats. This in turn could indicate increased psychomotor sensitization in the d-amphetamine-treated rats. Our data do not allow us to evaluate this possibility, as we did not measure stereotypy. Of note, if there was stereotypy, it was not strong enough to interfere with cocaine self-administration behaviour, as d-amphetamine-treated rats and control rats took similar amounts of cocaine (Fig. 4C). Whatever the case may be, d-amphetamine did not prevent the expression of psychomotor sensitization in previously cocaine-experienced rats, and this is at odds with previous findings in male rats (Allain et al., 2021). The neurobiological mechanisms underlying the development versus expression of psychomotor sensitization can be dissociable (Kalivas & Weber, 1988; Vezina & Stewart, 1990; Wolf, 1998). For example, drug effects in the ventral tegmental area preferentially mediate the development of sensitization, while drug effects in the nucleus accumbens preferentially mediate the expression of already established sensitization (Kalivas & Weber, 1988; Vezina & Stewart, 1990). One possibility is that in females, d-amphetamine more effectively disrupts cocaine effects that initiate the *development* of sensitization, while in males, d-amphetamine interferes with cocaine effects that underlie both the development and expression of sensitization.

Another difference in d-amphetamine’s effects between the sexes is that d-amphetamine reduced drug-primed reinstatement of cocaine seeking in males (Allain et al., 2021), but not in females here (Figs. 3C and 5K). However, this should be interpreted with caution, as we do not have a full dose-response curve for drug-primed reinstatement of cocaine seeking here.

### 4.3 Potential mechanisms underlying d-amphetamine’s therapeutic effects

Exposure to d-amphetamine might reduce the reinforcing effects of cocaine (Peltier et al., 1996) and/or promote cocaine’s aversive effects. If this were the case, one would expect d-amphetamine treatment to change ongoing cocaine self-administration behaviour. However, across two independent cohorts, d-amphetamine did not significantly change ongoing cocaine intake [see also (Chiodo et al., 2008)]. Another possibility is that our d-amphetamine dosing regimen is neurotoxic or debilitating. However, neurotoxic effects (as indicated by axon terminal degeneration or tyrosine hydroxylase immunoreactive patches) are seen at much higher doses than used here [> 16-20 mg/kg/day; (Ricaurte et al., 1984; Ryan et al., 1990, 1988)]. In addition, studies using a range of different d-amphetamine treatment regimens in non-human primates or rats show no or only transient effects on operant responding for food (Chiodo et al., 2008; Czoty et al., 2010, 2011; Negus & Mello, 2003). This argues against any non-specific or debilitation-related explanations. Because rats showed reduced incentive motivation for cocaine long after the termination of d-amphetamine treatment, we hypothesize that d-amphetamine reduces cocaine use by ‘inoculating’ against forms of cocaine-induced neuroplasticity that increase motivation for cocaine. In support, Siciliano et al. (2018) showed that the ability of d-amphetamine treatment to reduce the motivation to take cocaine in rats potentially involves reversal of tolerance to cocaine-induced dopamine uptake inhibition at the dopamine transporter (DAT) in the nucleus accumbens core. However, Long Access procedures were used. Long Access cocaine intake promotes tolerance to drug evoked dopamine activity, whereas intermittent cocaine intake sensitizes to cocaine’s effects on dopamine (Calipari et al., 2013; Kawa et al., 2019a,b; Samaha et al., 2021). Indeed, studies using *ex vivo* brain slices found that IntA cocaine intake increases cocaine-induced DAT inhibition (Long Access intake had the opposite effect) and increases electrically evoked dopamine release in the nucleus accumbens core (Calipari et al., 2013, 2014b). *In vivo* microdialysis studies show that IntA (but not Long Access) rats show sensitization of cocaine evoked dopamine overflow (Kawa et al., 2019b). Intermittent access cocaine experience also promotes psychomotor sensitization (Algallal et al., 2020; Allain et al., 2021, 2017b; Carr et al., 2020; Garcia et al., 2020), which is linked to enhanced dopamine transmission (Robinson & Berridge, 1993). Accordingly, d-amphetamine treatment might attenuate psychomotor and incentive sensitization to cocaine by preventing sensitization to cocaine’s dopamine elevating effects. In agreement, in male rats, d-amphetamine treatment prevents sensitization-related changes in cocaine potency at the DAT otherwise produced by IntA cocaine experience (Allain et al., 2021). We predict that similar mechanisms are at play in females, especially given that IntA cocaine intake is particularly effective in producing sensitization-related plasticity in female animals (Algallal et al., 2020; Carr et al., 2020; Kawa et al., 2019a). A previous study has examined the effects of d-amphetamine on striatal dopamine dynamics in female rats with a history of chronic cocaine self-administration (Siciliano et al., 2019). Cocaine self-administration was found to decrease both the rate of dopamine uptake and cocaine’s potency at the DAT in the nucleus accumbens core (decreased cocaine potency was also seen in the dorsolateral caudate-putamen). Notably, d-amphetamine exposure restored both effects. In accord with previous work (Allain et al., 2021; Siciliano et al., 2018), these findings highlight d-amphetamine’s ability to restore dopamine terminal function after chronic cocaine self-administration. However, the rats used in Siciliano et al. (2019) had Long Access rather than IntA to cocaine. Furthermore, d-amphetamine treatment was acute, rather than chronic. While chronic d-amphetamine exposure reduces cocaine use (Abreu, 1988; Allain et al., 2021; Chiodo et al., 2008; Chiodo & Roberts, 2009; Czoty et al., 2010, 2011; Grabowski et al., 2001, 2003; Greenwald et al., 2010; Lile et al., 2020; Morgane et al., 2013; Negus, 2003; Negus & Mello, 2003; Nuijten et al., 2016; Rush et al., 2010; Shearer et al., 2003; Siciliano et al., 2018; Zimmer et al., 2014), acute d-amphetamine administration can increase the response to cocaine (Barrett et al., 2004; Li et al., 2006) and also act as a prime to reinstate cocaine-seeking behaviour (Lynch et al., 1998; Schenk & Partridge, 1999). Thus, further work is needed to better understand the neurobiological basis of d-amphetamine effects on cocaine use in females and males. Of note, d-amphetamine treatment reduced cocaine-seeking under extinction conditions, in the absence of cocaine. As such, d-amphetamine likely reduces responding for cocaine via several neurobiological mechanisms, including but not limited to attenuation of sensitization-related changes at the DAT.

## 5. Conclusion

In female rats, d-amphetamine treatment during cocaine self-administration later reduced incentive motivation for cocaine, as indicated by decreased responding for the drug both under a PR schedule of reinforcement and under extinction conditions. We observed these effects in both cocaine-naïve and cocaine-experienced rats, thus increasing relevance for treatment of patients presenting with a history of chronic cocaine use. The therapeutic effects of d-amphetamine were also observed days after the discontinuation of treatment. Thus, efficacy long outlasted d-amphetamine’s presence in the body, suggesting persistent effectiveness, even without daily maintenance. Our findings are particularly relevant in the context of work showing that in treating drug addiction, best practices should include pharmacotherapy, in addition to cognitive behavioural therapy (Ray et al., 2020). Thus, our results will inform future work on the mechanisms underlying the potential therapeutic effects of d-amphetamine therapy in cocaine addiction across the sexes.

## Acknowledgments

We thank Ana Hernández Sauret for technical assistance with behavioural testing. We also thank Dr Mike J.F. Robinson for comments on an earlier version of this text.

## Author contributions

NAN and ANS designed the research. NAN conducted all experiments with guidance from FA and ANS. NAN analysed data with guidance from FA and ANS. NAN and ANS wrote the manuscript with revisions from FA.

---

**Figure S1.**
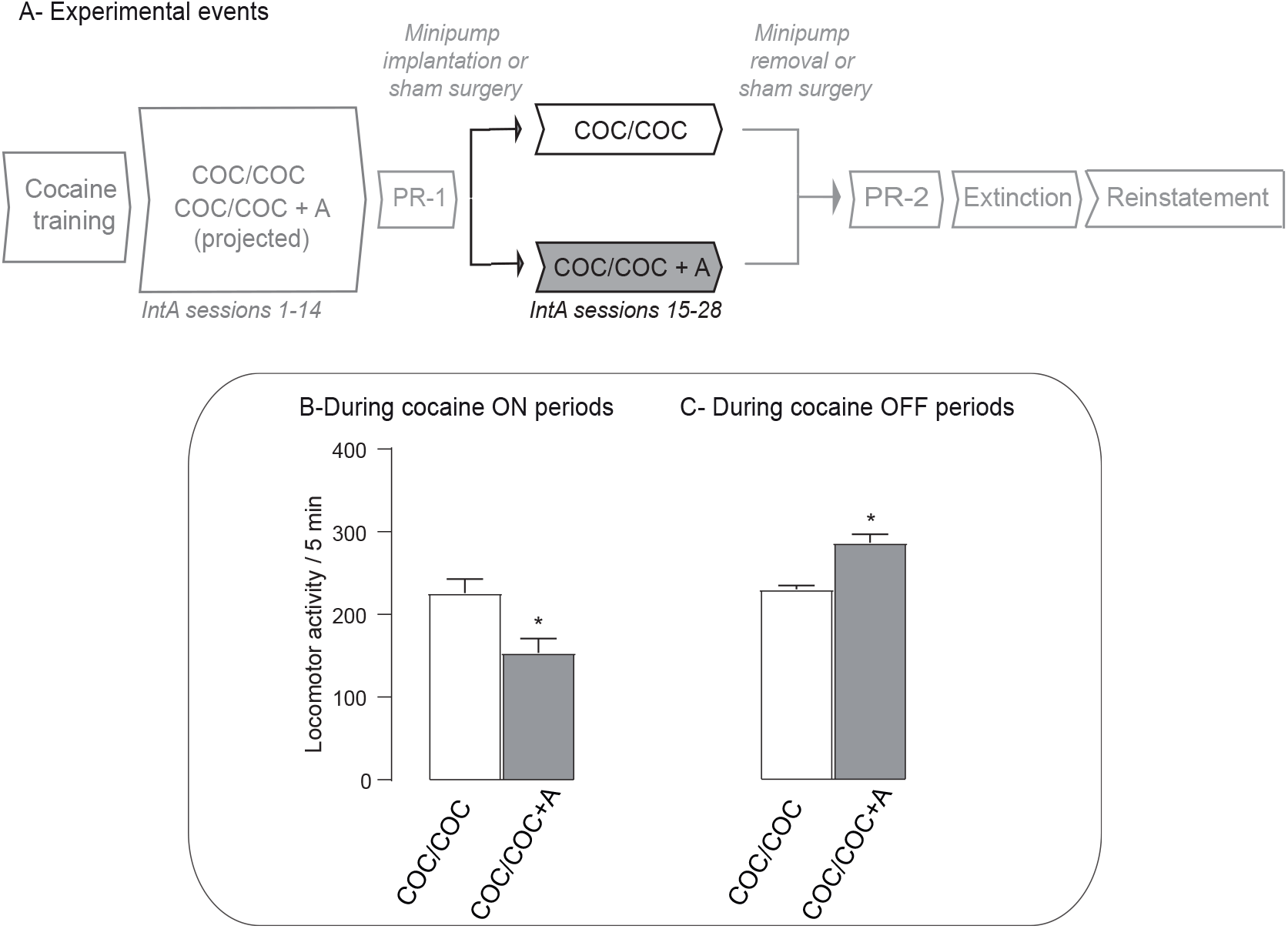
In cocaine-experienced rats, D-amphetamine treatment during intermittent cocaine self-administration sessions decreased locomotor activity during cocaine ON periods, and increased locomotor activity during cocaine OFF periods. (A) Sequence of experimental events for experiment 2. (B), d-amphetamine-treated rats showed reduced locomotor activity during the 5-minute cocaine ON periods of the last Intermittent-access session (i.e., session 28). (C), d-amphetamine-treated rats showed enhanced locomotor activity during the 5-minute cocaine OFF periods of the last Intermittent-access session. Data are mean ± SEM. n = 7 for the COC/COC group and n = 9 for the COC/COC + A group. * p < 0.0001.

